## Supplemental material for "Marcks and Marcks-like 1 proteins promote spinal cord development and regeneration in *Xenopus*"

**Supplementary figures**

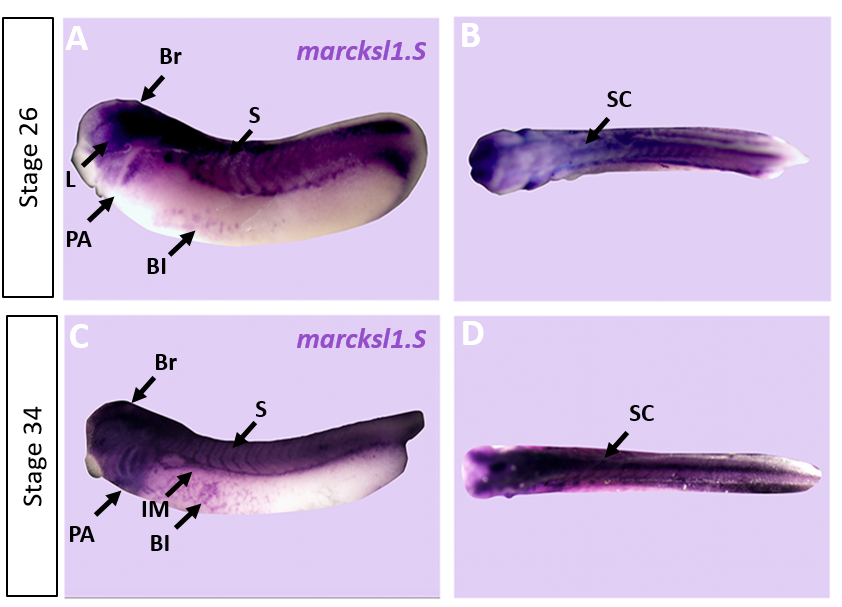

**Fig. S1: Expression of *marcksl1. S* in *Xenopus.***

Expression of *marcksl1.S* as revealed by whole mount in situ hybridization in stage 26 (A-B) and 34 (C-D) *Xenopus* embryos in lateral (A,C) and dorsal (B, D) views (anterior is to the left). Expression in the brain (Br), spinal cord (SC), lens (L), somites (S), intermediate mesoderm (IM), blood islands (Bl), and pharyngeal arches (PA) is indicated.

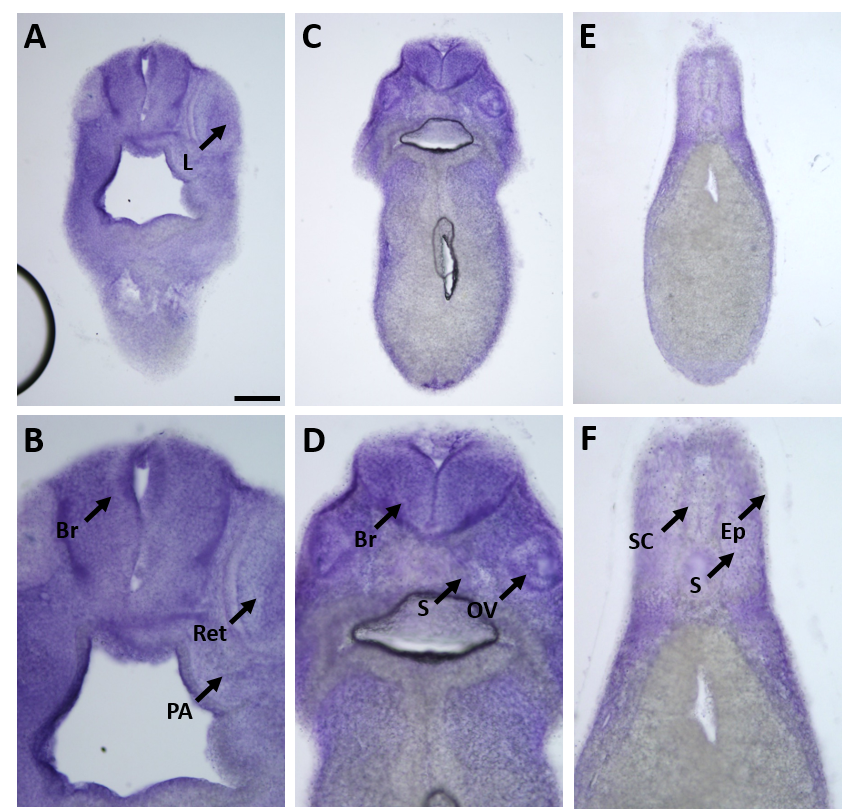

**Fig. S2: Expression of *marcks*.*S* mRNA revealed by transverse vibratome sections of *Xenopus* stage 34 embryos after whole-mount in situ hybridization.**

Higher magnification details of (A, C, E) are shown in (B, D, F), respectively. *marcks*.S expression is evident in the brain (Br), retinas (Ret), lens (L), spinal cord (SC), PA, otic vesicle (OV), epidermis (Ep), and somites (S). Scale bar in A = 100 µm for (A, C, E) and 50 µm for (B, D, F).

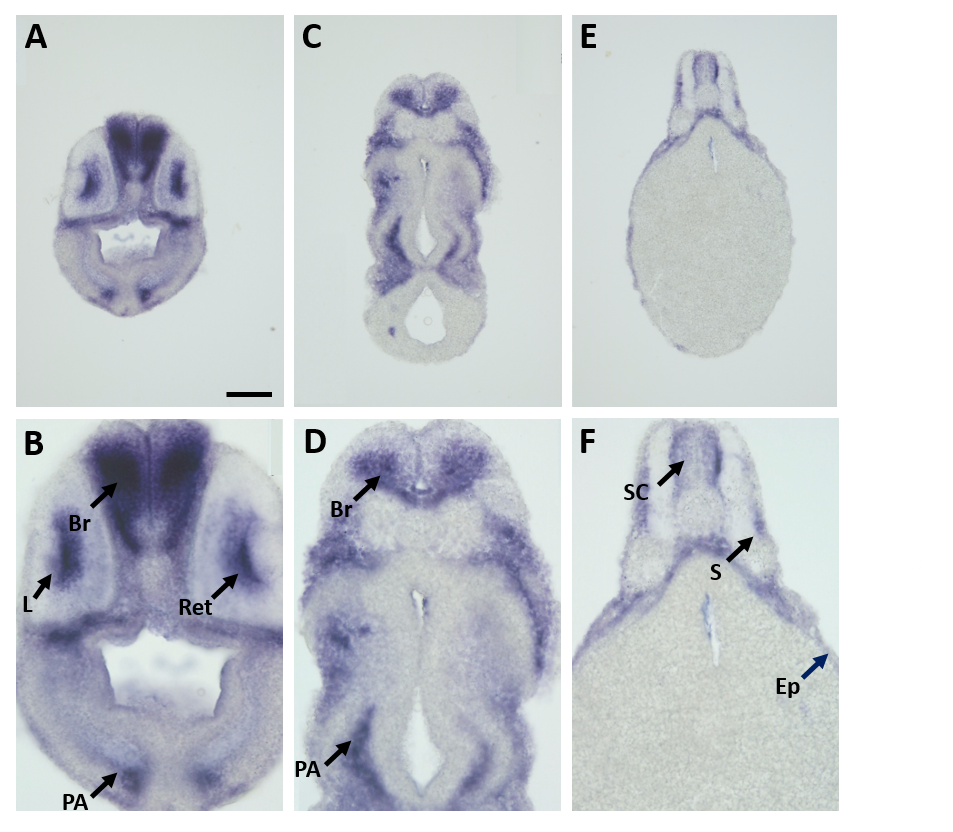

**Fig. S3: Expression of *marcksl1.L* mRNA as revealed by transverse vibratome sections of *Xenopus* stage 34 embryos after whole-mount in situ hybridization.**

Higher magnification details of (A, C, E) are shown in (B, D, F), respectively. *marcks*l1.L expression is evident in the brain (Br), retinas (Ret), lens (L), spinal cord (SC), PA, epidermis (Ep), and somites (S). Scale bar in A = 100 µm for (A, C, E) and 50 µm for (B, D, F).

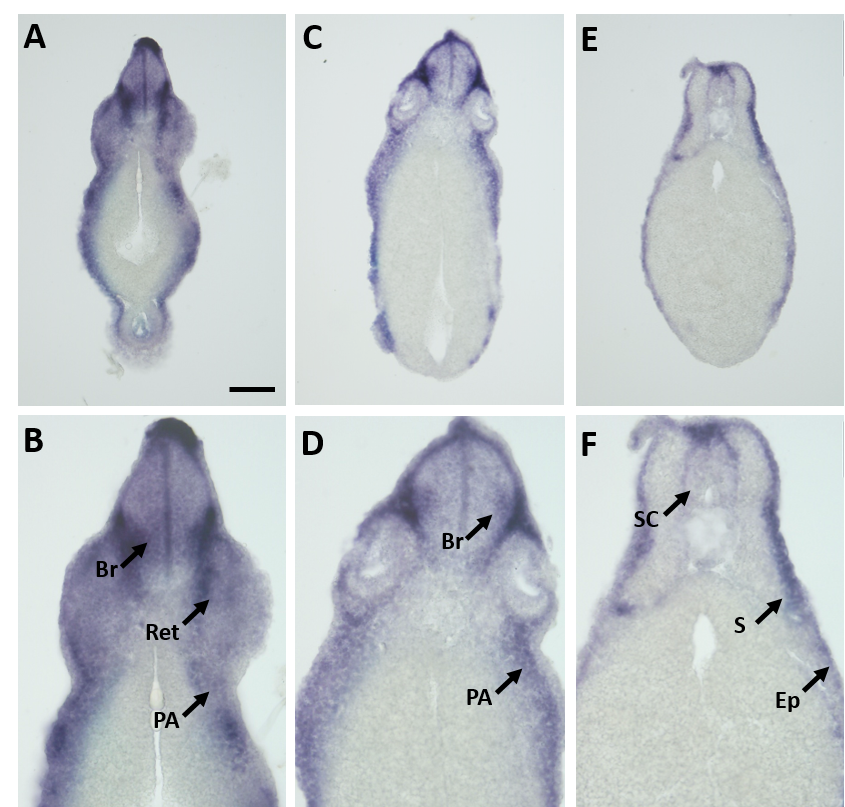

**Fig. S4: Expression of *marcksl1.S* mRNA as revealed by transverse vibratome sections of *Xenopus* stage 34 embryos after whole-mount in situ hybridization.**

Higher magnification details of (A, C) are shown in (B, D). *marcks*l1.L expression is evident in the brain (Br), retinas (Ret), spinal cord (SC), pharyngeal arches (PA), epidermis (Ep), and somites (S). Scale bar in A = 100 µm for (A, C) and 50 µm for (B, D).

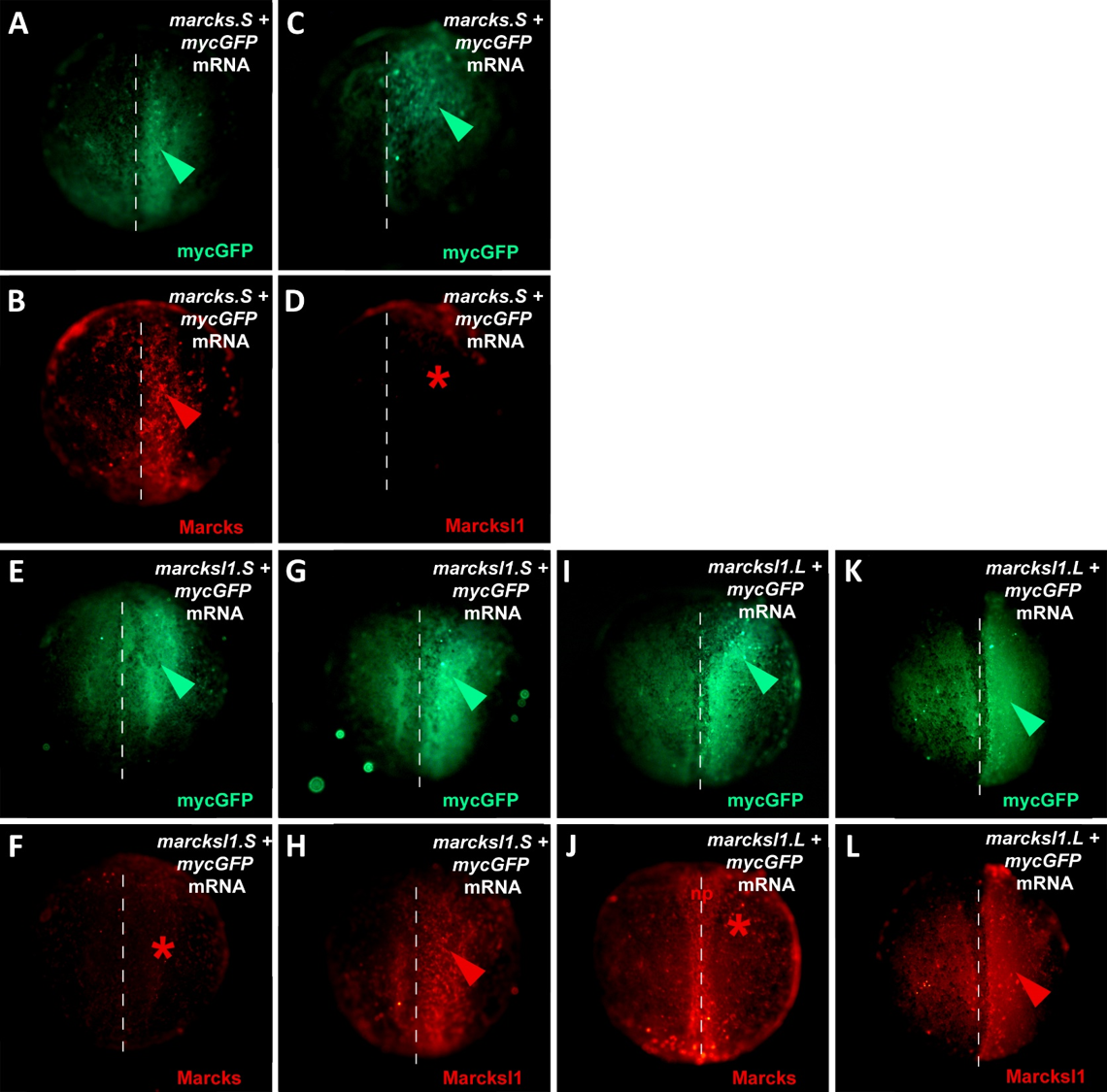

**Fig. S5: Specificity of Marcks and Marcksl1 antibodies.**

One blastomere of 2-cell stage embryos was co-injected with 125 pg of mycGFP mRNA and 500 pg of either *marcksS* mRNA (A-D), *marcksl1.S* mRNA (E-H), or *marcksl1.L* mRNA (I-L) followed by immunostaining at neural plate stages (stage 13-16) with anti-Myc (A, C, E, G, I, K) and either anti-Marcks (B, F, J) or anti-Marcksl1 (D, H, L) antibodies. The injected side is shown on the right, with the injected area indicated by mycGFP immunostaining (green arrowheads). Anti-Marcks antibodies recognize Marcks (B, red arrowhead) but not Marcksl1 (F, I, red asterisks) proteins, while anti-Marcksl1 antibodies recognize Marcksl1 (H, L, red arrowheads) but not Marcks (D, red asterisk) proteins translated from the injected mRNAs indicating that antibodies do not cross-react. Marcks antibodies also detected native Marcks1 protein in the neural plate in (np) in embryos at late neural plate stages (J).

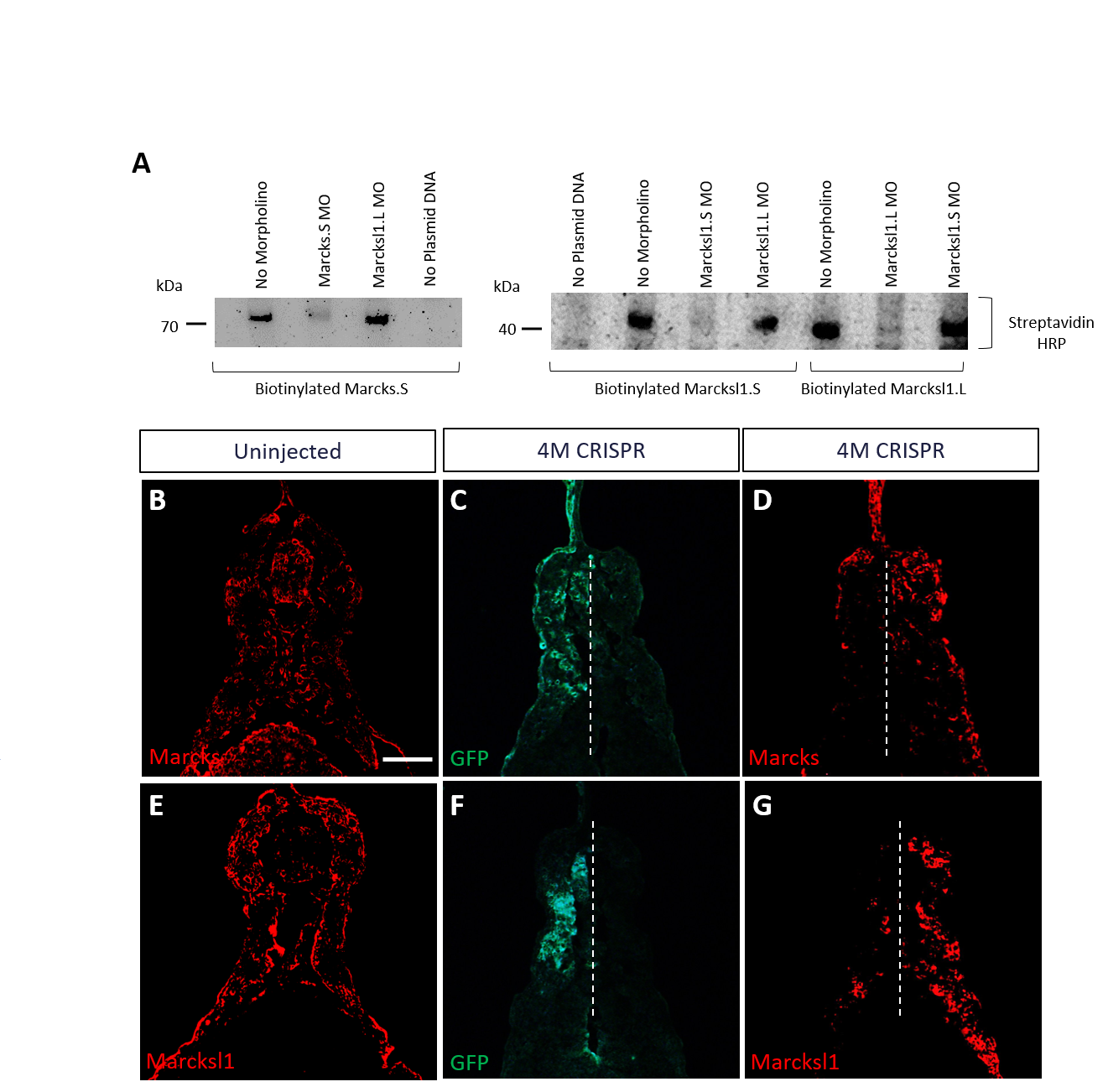

**Fig. S6: Validation of Marcks and Marcksl1 knockdown and knockout.**

A: Morpholino (MO) specificity and efficacy verified by in vitro TNT. Biotinylated Marcks.S, Marcksl1.L, and Marcksl1.S proteins were expressed by in vitro transcription and translation (TNT) and detected in western plots by Streptavidin-HRP. In vitro, TNT reactions without any morpholino added served as positive control. In vitro, TNT reactions without plasmid DNA added served as negative control. Protein translation was blocked when the appropriate MO was added, but not by an inappropriate MO. B-G: Reduction of Marcks and Marcksl1 immunostaining in stage 36 embryos co-injected with *marcks.L/S*, *marcksl1.L/S* sgRNA and *myc-GFP* mRNA (4M CRISPR). In uninjected embryos (B, E), Marcks (B) and Marcksl1 (E) expression are seen on both sides. In 4M CRISPR-injected embryos Marcks (D) and Marcksl1 (G) immunostaining is reduced on the injected side as indicated by GFP immunostaining (C, F). Scale bar in B = 50 µm (for all panels).

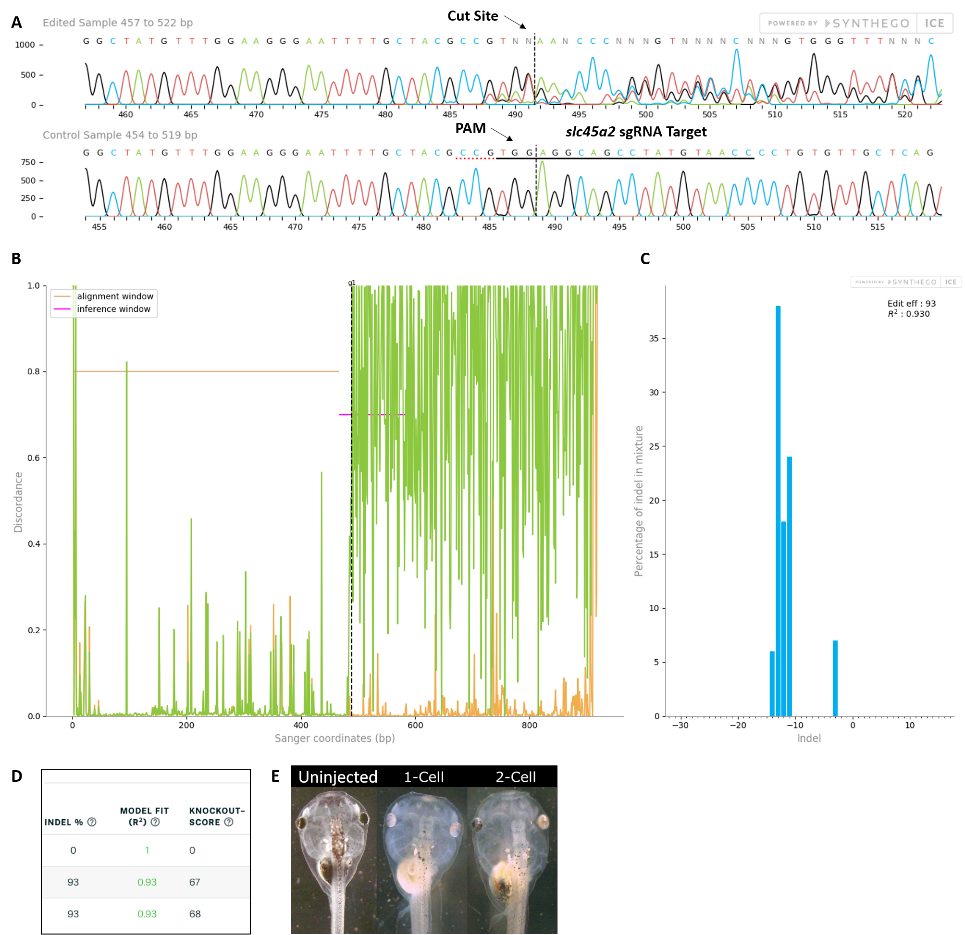

**Fig. S7: Control *slc45a2* sgRNA/Cas9 efficiently and specifically edits targeted *X. laevis* DNA.**

All data were generated by the Synthego Inference for CRISPR Edits (ICE) tool based on DNA samples from stage 26-42 embryos injected with slc45a2 sgRNA, Cas9 protein, and *myc-GFP* mRNA during the 1-cell stage in comparison with uninjected embryos. A: Sanger sequence chromatograms showing degradation of the sequence trace from PCR products amplifying CRISPR-targeted *slc45a2* DNA (top) compared with DNA from uninjected embryos (bottom). The underlined region indicates the sgRNA target binding sequence. Arrows indicate the Protospacer Adjacent Motif (PAM) and the Cas9 cleavage site. B: Discordance plot showing the level of sequence-base disagreement between the control reference DNA from uninjected embryos (orange lines) and the CRISPR DNA (green lines) in the region surrounding the cut site, indicating a high level of sequence difference after the cleavage site, shown by the hatched line. C: Synthego Indel Distribution plot, which infers the distribution of insertion-deletion sizes (± 1 or more nucleotides) around the *slc45a2* CRISPR-target site. The x-axis indicates the size of the indel, and the y-axis shows the percentage of the sequences containing it. The editing efficiency score indicates that 93% of the DNA in the CRISPR-targeted sequence was edited. A Pearson correlation coefficient (R^2^) was also generated by ICE with a value of 0.93 out of 1, measuring how well the indel distribution fits the Sanger sequence data. D: Summary of three analyzed samples showing the sgRNA target and PAM sequences. DNA from PCR products comparing two *slc45a2* sequences from uninjected embryos (top) shows an indel percentage of 0% and a predicted knockout (KO) score of 0%. Indel scores for DNA sequences derived from two CRISPR-edited embryos (middle and bottom) indicate that 93% of the DNA analyzed was edited. Knockout scores of 67-68% indicate the proportion of indels that contain a frameshift mutation or a 21+ base-pair indel. E: *slc45a2* CRISPR knockout generates a visible loss of eye pigmentation in stage 46 tadpoles. Whereas uninjected tadpoles have no eye pigmentation defects, eye pigmentation is unilaterally reduced in the CRISPR-injected side of 2-cell stage-injected embryos and bilaterally reduced in 1-cell stage CRISPR-injected embryos.

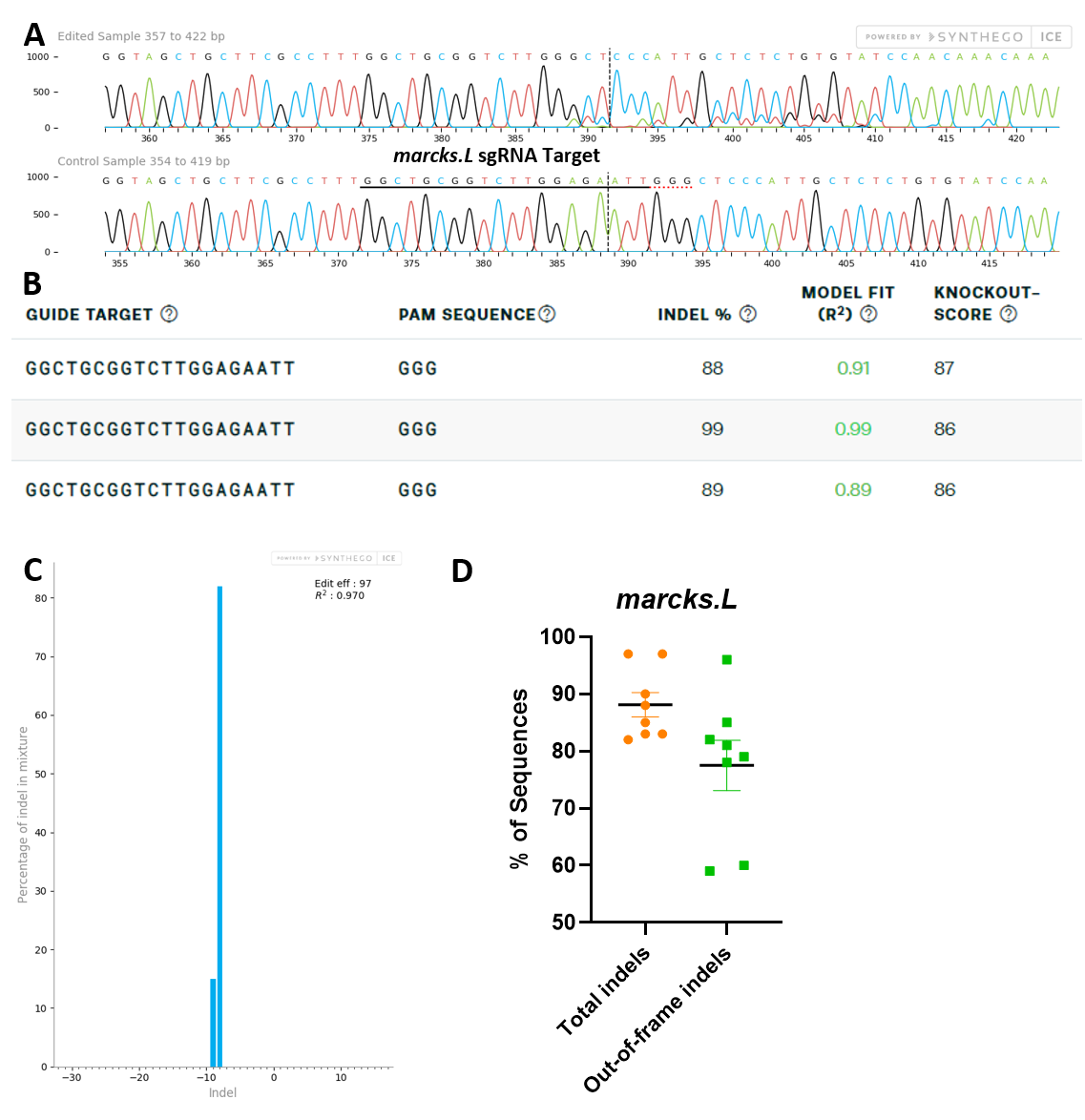

**Fig. S8:** ***marcks.L/S* sgRNA efficiently and specifically edits *Xenopus* *laevis* *marcks.L* DNA.**

All data were generated by the Synthego Inference for CRISPR Edits (ICE) tool based on DNA samples from stage 26-42 embryos injected with marcks.L/S and marcksl1.L/S sgRNA, Cas9 protein, and *myc-GFP* mRNA during the 1-cell stage in comparison with uninjected embryos. A: Sanger sequence chromatograms showing degradation of the sequence trace from PCR products amplifying CRISPR-targeted *marcks.L* exon 1 DNA (top) compared with DNA from an uninjected embryo (bottom). The underlined region indicates the sgRNA target binding sequence. B: Summary of three analyzed samples showing the sgRNA target and PAM sequence. Indel scores for DNA sequences derived from 3 CRISPR-edited embryos compared with uninjected embryos, indicating that 88%, 89%, and 99% of the DNA analyzed was edited. Knockout scores of 86% and 87% indicate the proportion of indels that contain a frameshift mutation. C: Synthego Indel Distribution plot from one representative CRISPR-embryo, which infers the distribution of insertion-deletion sizes (± 1 or more nucleotides) around the *marcks.L* CRISPR-target site. The x-axis indicates the size of the indel, and the y-axis shows the percentage of the sequences containing it. The editing efficiency score indicates that 97% of the DNA in the CRISPR-targeted sequence was edited with mostly an eight-base-pair deletion. A Pearson correlation coefficient (R^2^) was also generated by ICE with a value of 0.97 out of 1, measuring how well the indel distribution fits the Sanger sequence data. D: Indel percentages calculated by ICE for 8 CRISPR-edited embryos, indicating that an average of 88% of the DNA was edited for all animals. From this edited DNA, 78% contained out-of-frame mutations.

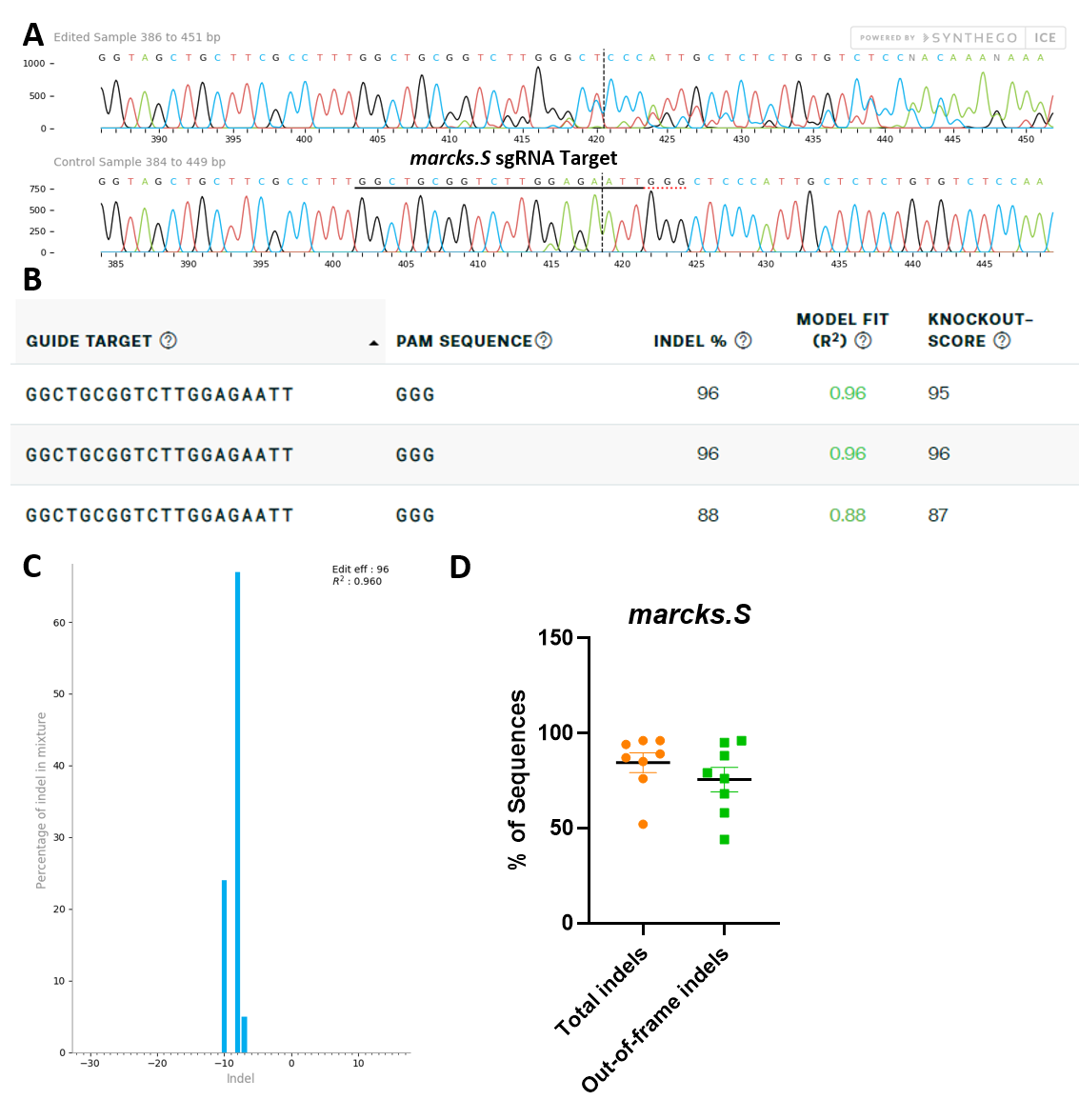

**Fig. S9:** ***marcks.L/S* sgRNA efficiently and specifically edits *Xenopus* *laevis* *marcks.S* DNA.**

All data were generated by the Synthego ICE tool based on DNA samples from stage 26-42 embryos injected with marcks.L/S and marcksl1.L/S sgRNA, Cas9 protein, and *myc-GFP* mRNA during the 1-cell stage in comparison with uninjected embryos. A: Sanger sequence chromatograms showing degradation of the sequence trace from PCR products amplifying CRISPR-targeted *marcks.S* exon 1 DNA (top) compared with DNA from an uninjected embryo (bottom). The underlined region indicates the sgRNA target binding sequence. B: Summary of three analyzed samples showing the sgRNA target and PAM sequence. Indel scores for DNA sequences derived from 3 CRISPR-edited embryos compared with uninjected embryos, indicating that 88% and 96% of the analyzed DNA was edited. Knockout scores of 95%, 96%, and 87% indicate the proportion of indels that contain a frameshift mutation. C: Synthego Indel Distribution plot from one representative CRISPR-embryo, which infers the distribution of insertion-deletion sizes (± 1 or more nucleotides) around the *marcks.S* CRISPR-target site. The x-axis indicates the size of the indel, and the y-axis shows the percentage of the sequences containing it. The editing efficiency score indicates that 96% of the DNA in the CRISPR-targeted sequence was edited with a majority of eight base-pair deletions. A Pearson correlation coefficient (R^2^) was also generated by ICE with a value of 0.96 out of 1, measuring how well the indel distribution fits the Sanger sequence data. D: Indel percentages calculated by ICE for 8 CRISPR-edited embryos, indicating that an average of 84% of the DNA was edited for all animals. From this edited DNA, 76% contained out-of-frame mutations.

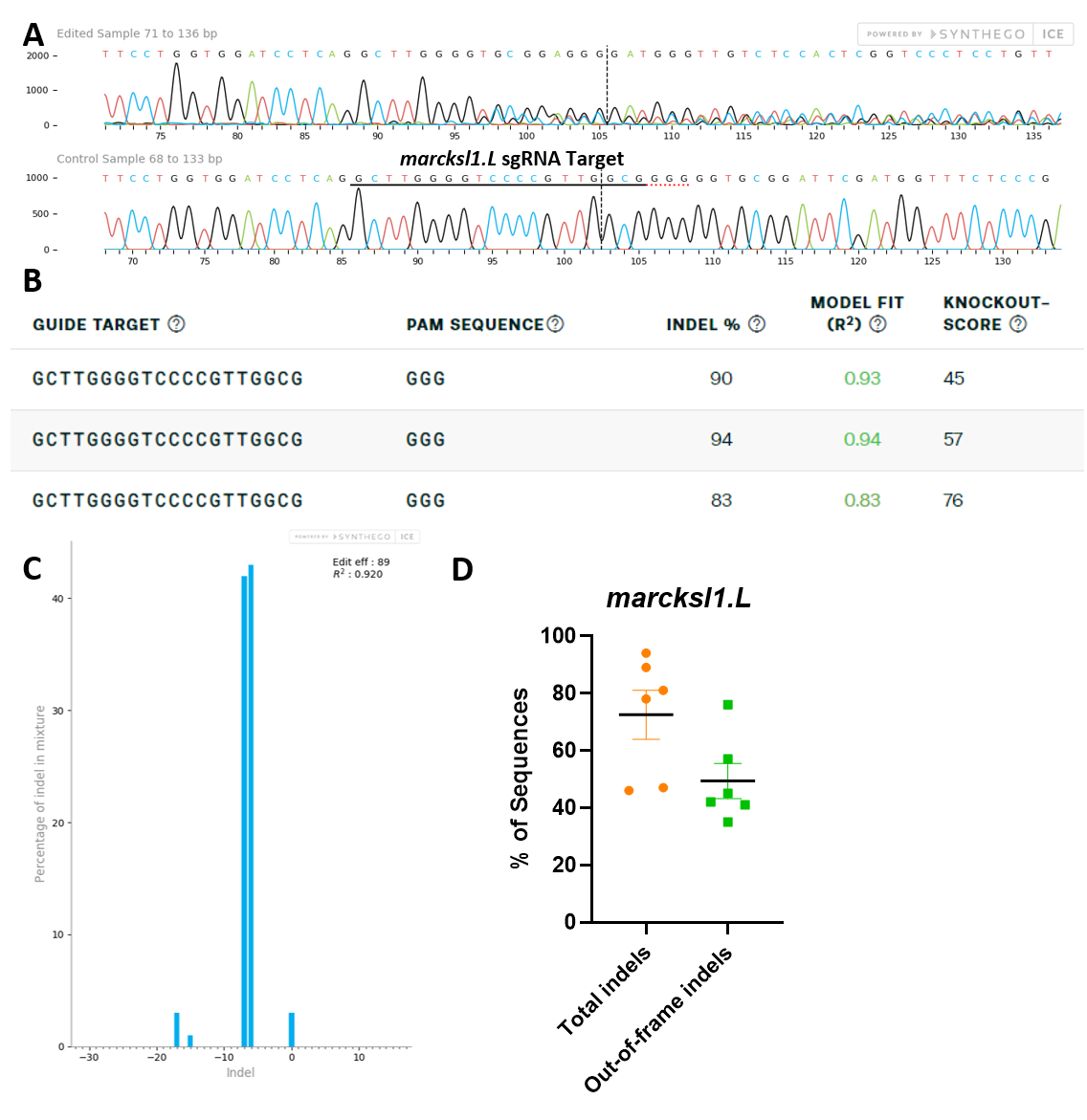

**Fig. S10: *marcksl1.L/S* sgRNA efficiently and specifically edits *X.* *laevis* *marcksl1.L* DNA.**

All data were generated by the Synthego Inference for CRISPR Edits (ICE) tool based on DNA samples from stage 26-42 embryos injected with marcks.L/S and marcksl1.L/S sgRNA, Cas9 protein, and *myc-GFP* mRNA during the 1-cell stage in comparison with uninjected embryos. A: Sanger sequence chromatograms showing degradation of the sequence trace from PCR products amplifying CRISPR-targeted *marcksl1.L* exon 2 DNA (top) compared with DNA from an uninjected embryo (bottom). The underlined region indicates the sgRNA target binding sequence. B: Summary of three analyzed samples showing the sgRNA target and PAM sequence. Indel scores for DNA sequences derived from 3 CRISPR-edited embryos compared with uninjected embryos, indicating that 83%, 90%, and 94% of the DNA analyzed was edited. Knockout scores of 45%, 57%, and 76% indicate the proportion of indels that contain a frameshift mutation. C: Synthego Indel Distribution plot from one representative CRISPR-embryo, which infers the distribution of insertion-deletion sizes (± 1 or more nucleotides) around the *marcksl1.L* CRISPR-target site. The x-axis indicates the size of the indel, and the y-axis shows the percentage of the sequences containing it. The editing efficiency score indicates that 89% of the DNA in the CRISPR-targeted sequence was edited with mostly 6-7 base-pair deletions. A Pearson correlation coefficient (R^2^) was also generated by ICE with a value of 0.92 out of 1, measuring how well the indel distribution fits the Sanger sequence data. D: Indel percentages calculated by ICE for 6 CRISPR-edited embryos, indicating that an average of 73% of the DNA was edited for all animals. From this edited DNA, 50% contained out-of-frame mutations.

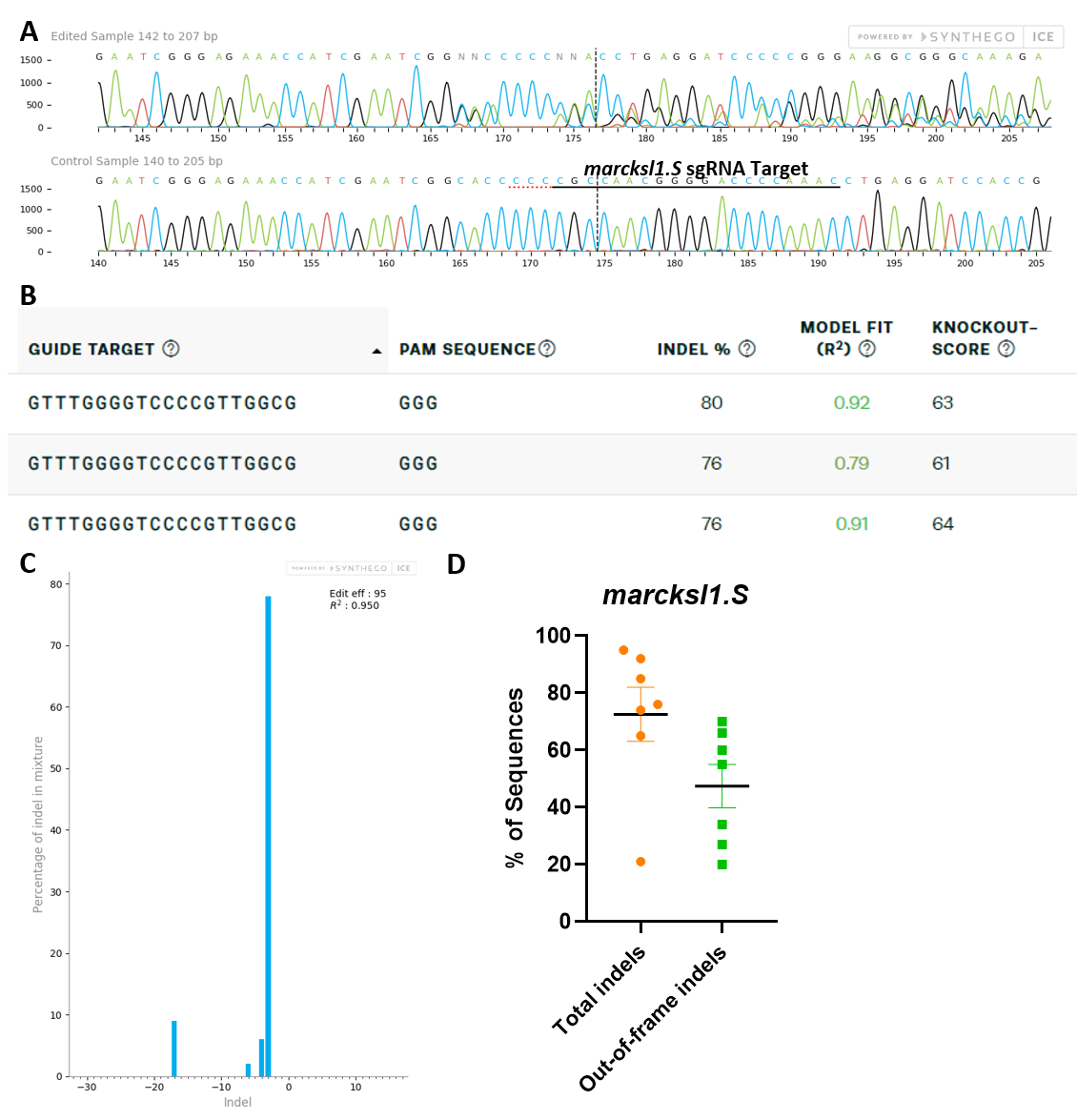

**Fig. S11: *marcksl1.L/S* sgRNA efficiently and specifically edits *X.* *laevis* *marcksl1.S* DNA.**

All data were generated by the Synthego Inference for CRISPR Edits (ICE) tool based on DNA samples from stage 26-42 embryos injected with marcks.L/S and marcksl1.L/S sgRNA, Cas9 protein, and *myc-GFP* mRNA during the 1-cell stage in comparison with uninjected embryos. A: Sanger sequence chromatograms showing degradation of the sequence trace from PCR products amplifying CRISPR-targeted *marcksl1.S* exon 2 DNA (top) compared with DNA from an uninjected embryo (bottom). The underlined region indicates the sgRNA target binding sequence. B: Summary of three analyzed samples showing the sgRNA target and PAM sequence. Indel scores for DNA sequences derived from 3 CRISPR-edited embryos compared with uninjected embryos, indicating that 76% and 80% of the DNA analyzed was edited. Knockout scores of 61%, 63%, and 64% indicate the proportion of indels that contain a frameshift mutation. C: Synthego Indel Distribution plot from one representative CRISPR-embryo, which infers the distribution of insertion-deletion sizes (± 1 or more nucleotides) around the *marcksl1.S* CRISPR-target site. The x-axis indicates the size of the indel, and the y-axis shows the percentage of the sequences containing it. The editing efficiency score indicates that 95% of the DNA in the CRISPR-targeted sequence was edited with mostly a 3-base-pair deletion. A Pearson correlation coefficient (R^2^) was also generated by ICE with a value of 0.95 out of 1, measuring how well the indel distribution fits the Sanger sequence data. D: Indel percentages calculated by ICE for 7 CRISPR-edited embryos, indicating that an average of 73% of the DNA was edited for all animals. From this edited DNA, 47% contained out-of-frame mutations.

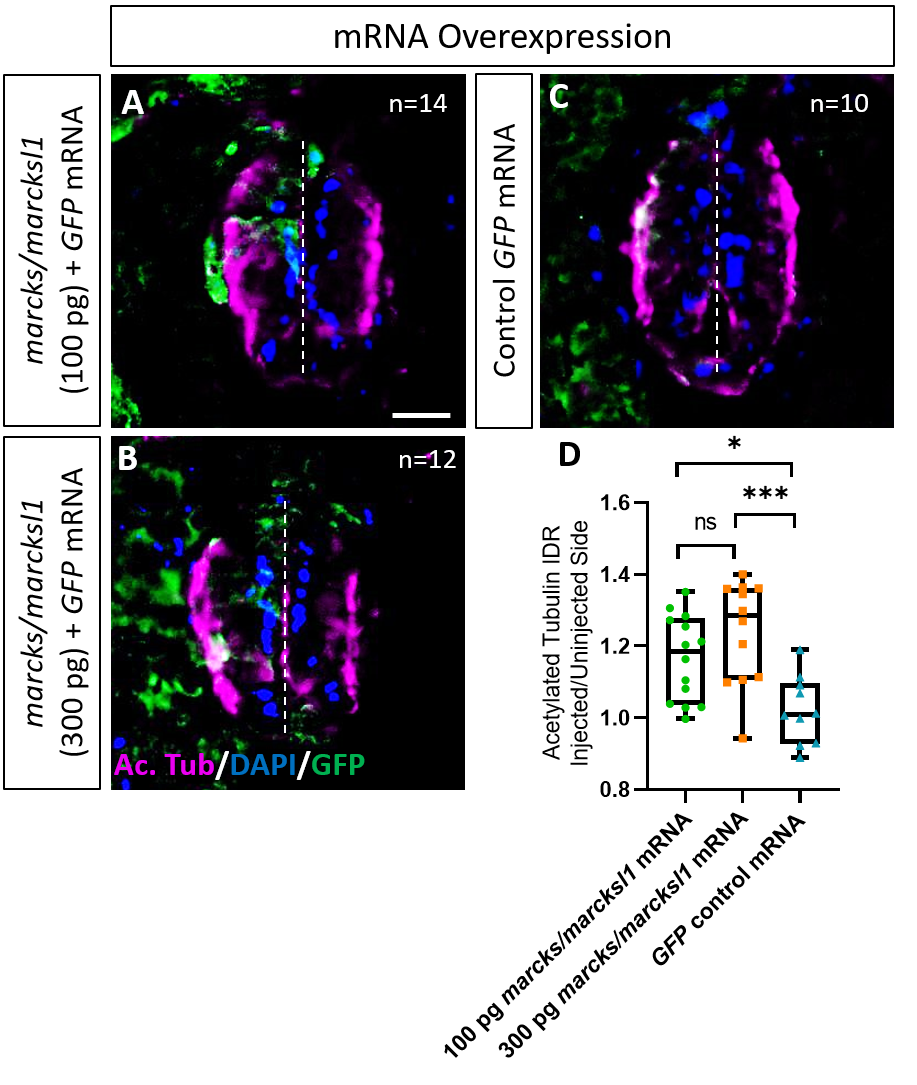

**Fig. S12:** **Marcks and Marcksl1 overexpression promotes neurite outgrowth during *Xenopus* spinal cord development.**

One blastomere of 4-8 cell stage embryos was co-injected with 100 pg (A) and 300 pg (B) of each *marcks.*S, *marcksl1.L,* and *marcksl1.S* mRNA in addition to 125 pg *myc-GFP* mRNA. In control embryos, 125 pg *myc-GFP* mRNA was injected (C). Injected (left) and uninjected (right) spinal cord sides of stage 34-36 embryos (separated by the dashed line) were compared in transverse sections immunostained for acetylated tubulin (A-C) and for GFP to determine the injected side. Compared to control embryos, immunostaining for acetylated tubulin was quantified (D) by determining the integrated density ratio (IDR) of staining in injected to uninjected sides. Significance was determined using an ordinary one-way ANOVA with Tukey’s multiple comparisons test. NS, not significant; *P<0.05; ***P<0.001. Error bars represent minimum to maximum values showing all points with the line at mean. n refers to the number of animals analyzed. Scale bar in A = 25 µm (for all panels).

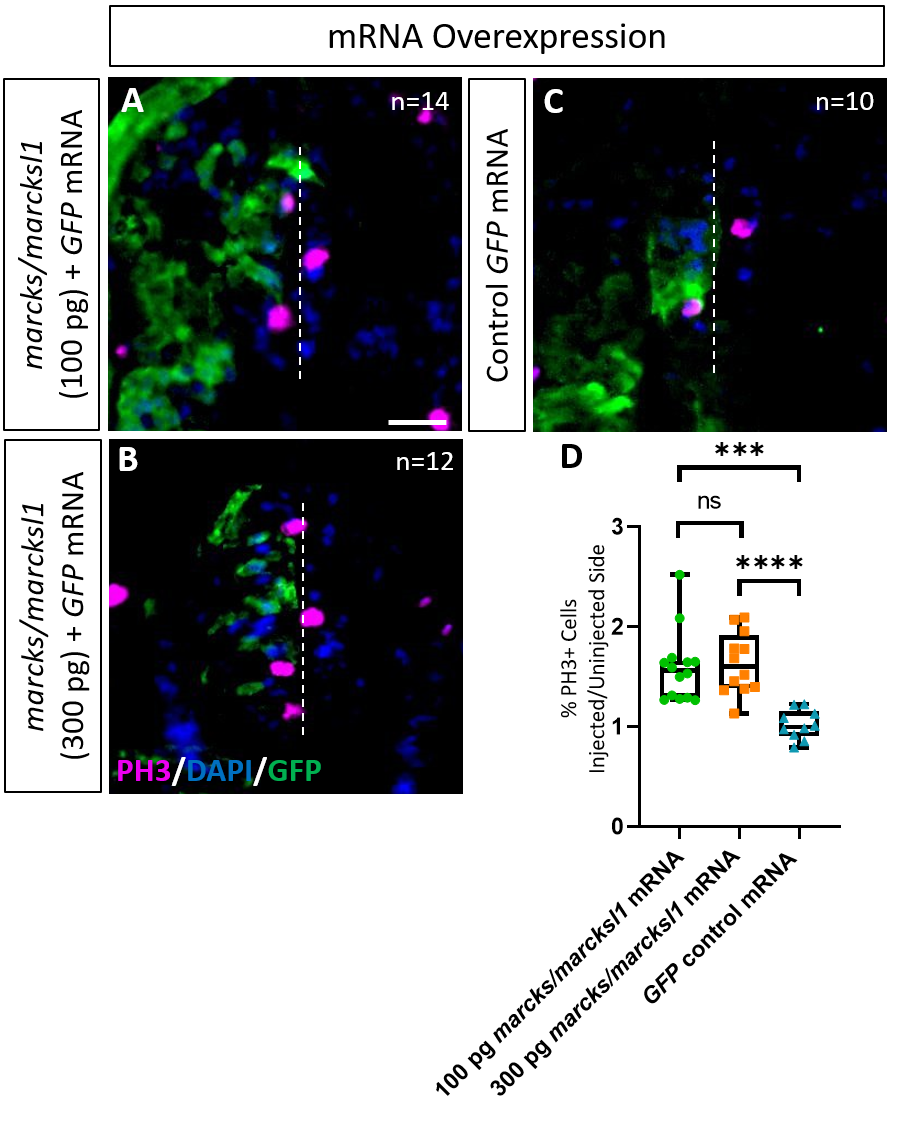

**Fig. S13: Marcks and Marcksl1 overexpression promotes mitotic activity during *Xenopus* spinal cord development.**

One blastomere of 4-8 cell stage embryos was co-injected with 100 pg (A) and 300 pg (B) of each *marcks.*S, marcksl1*.L,* and *marcksl1.S* mRNA in addition to 125 pg *myc-GFP* mRNA. In control embryos, 125 pg *myc-GFP* mRNA was injected (C). Injected (left) and uninjected (right) spinal cord sides of stage 34-36 embryos (separated by the dashed line) were compared in transverse sections immunostained for PH3 (A-C) and for GFP to determine the injected side. Immunostaining for acetylated tubulin was quantified (D) by determining the ratio of the percentage of PH3+ cells in injected and uninjected sides in comparison with control embryos. Significance was determined using an ordinary one-way ANOVA with Tukey’s multiple comparisons test. NS, not significant; ***P<0.001. Error bars represent minimum to maximum values showing all points with the line at the mean. n refers to the number of animals analyzed. Scale bar in A = 25 µm (for all panels).

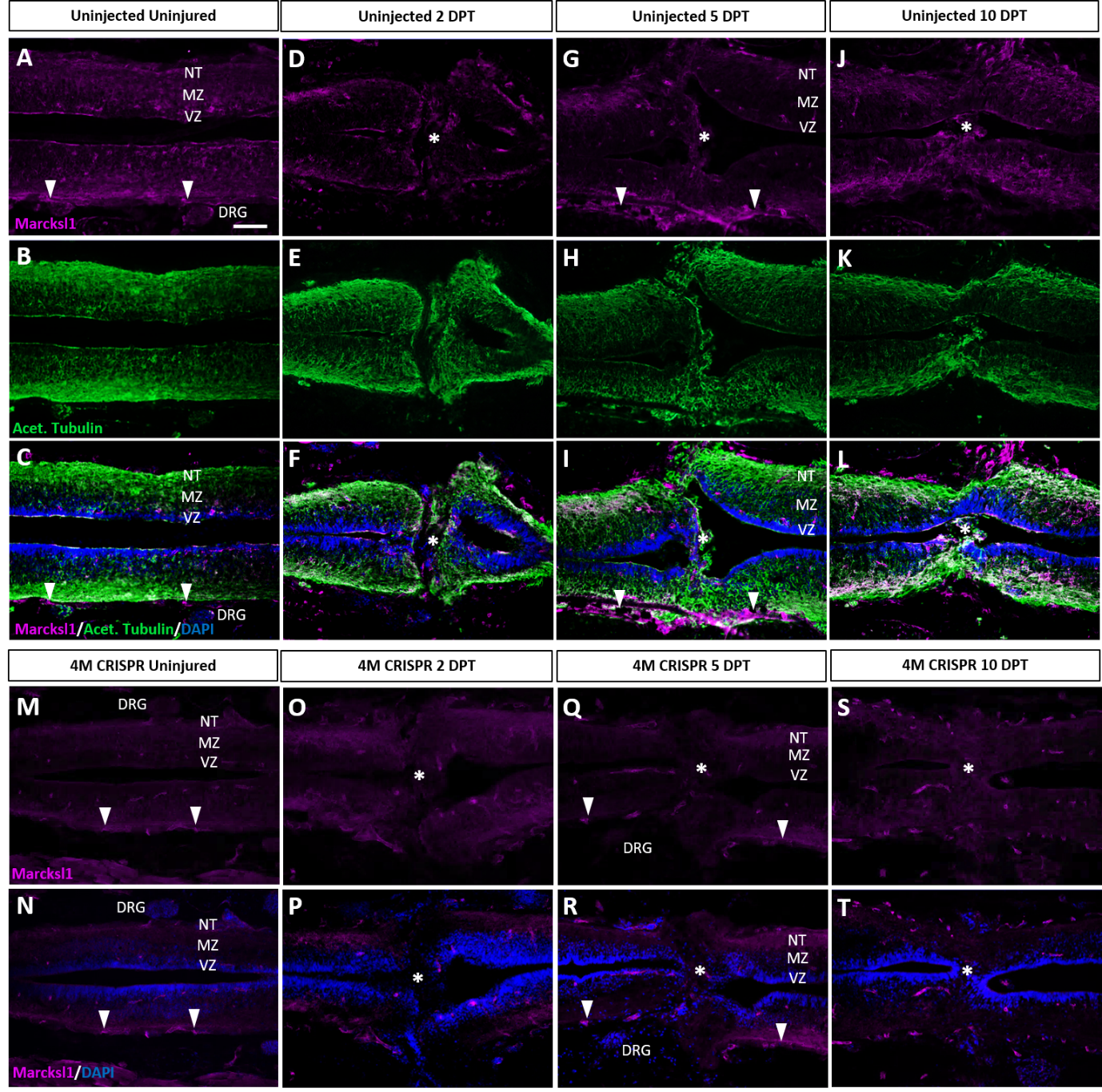

**Fig. S14 Marcksl1 proteins are upregulated after spinal cord transection in uninjected but not in 4M CRISPR-injected tadpoles.**

A-L: Immunostaining for Marcksl1 in horizontal sections of uninjected stage 50 tadpoles without injury (A-C) or at 2 DPT (D-F), 5 DPT (G-I), and 10 DPT (J-L). Before the injury, Marcksl1 immunostaining was weakly detected in the ventricular zone (VZ), mantle zone (MZ) and the neurite tracts (NT) of the marginal zone of the spinal as well as in the meninges (white arrowheads) and dorsal root ganglia (DRG). After spinal cord transection, it was slightly upregulated in cells infiltrating the injury site (white asterisks) and in all layers of the spinal cord surrounding the injury gap at 10 DPT. N=6 tadpoles per condition. M-T: Immunostaining for Marcksl1 in horizontal sections of 4M CRISPR-injected stage 50 tadpoles without injury (M, N) or at 2 DPT (O, P), 5 DPT (Q, R), and 10 DPT (S, T). Barely any Marcksl1 immunostaining is detectable at any time point. N=6 tadpoles per condition. Scale bar in A = 50 µm (for all panels).

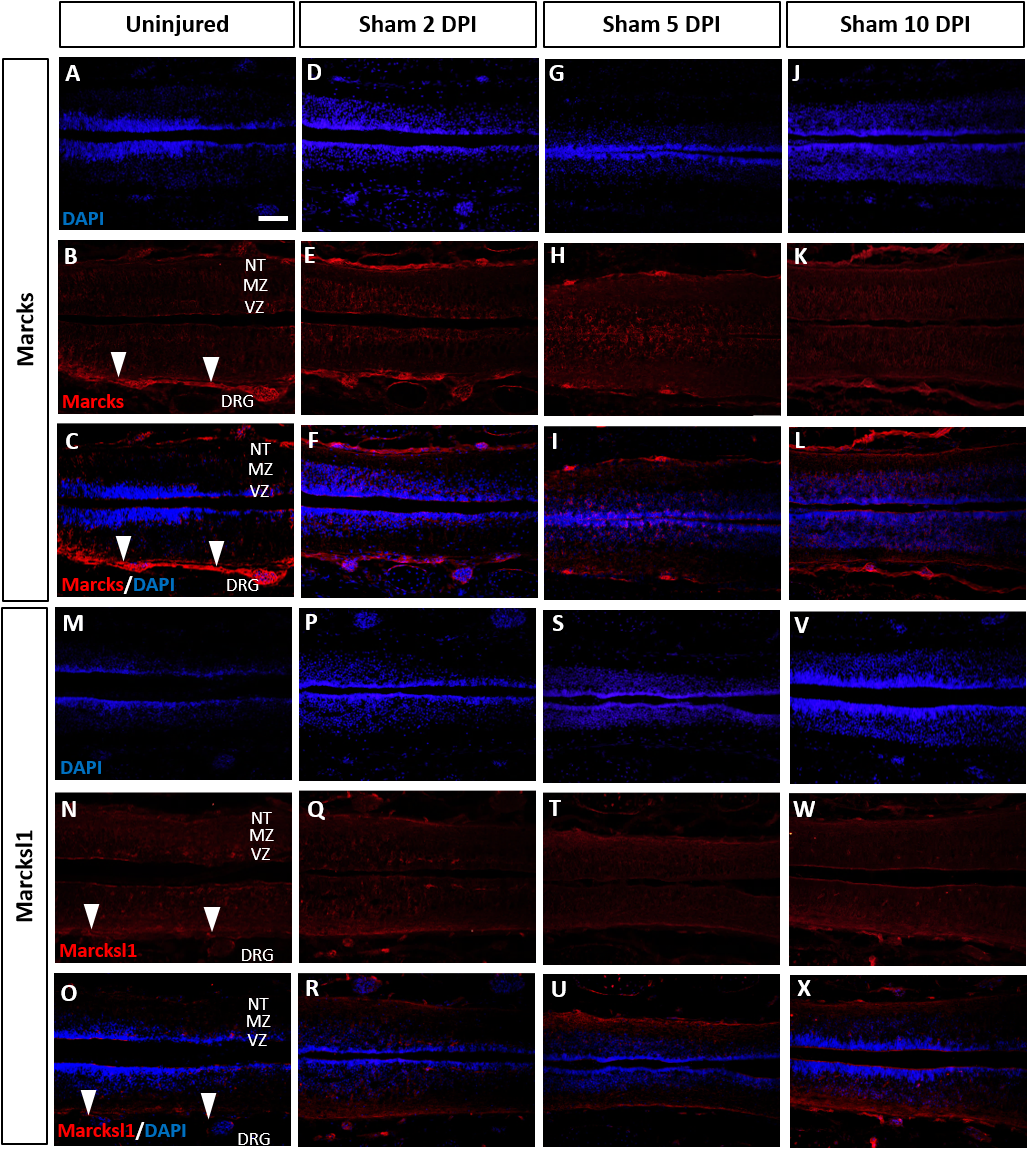

**Fig. S15: Marcks and Marcksl1 proteins do not change in uninjected sham-operated tadpoles.**

Immunostaining for Marcks (A-L) and Marcksl1 (M-X) in sham-operated tadpoles remained unaltered compared to the uninjured condition (A-C, M-O) at two days (D-F, P-R), five days (G-I, S-U), and ten days (J-L, V-X) post-injury (DPI). N= 3 tadpoles per condition. White arrowheads indicate the meninges. Abbreviations: DRG: dorsal root ganglia; MZ: mantle zone; NT of the marginal zone; VZ: ventricular zone. Scale bar in A= 50 µm (for all panels).

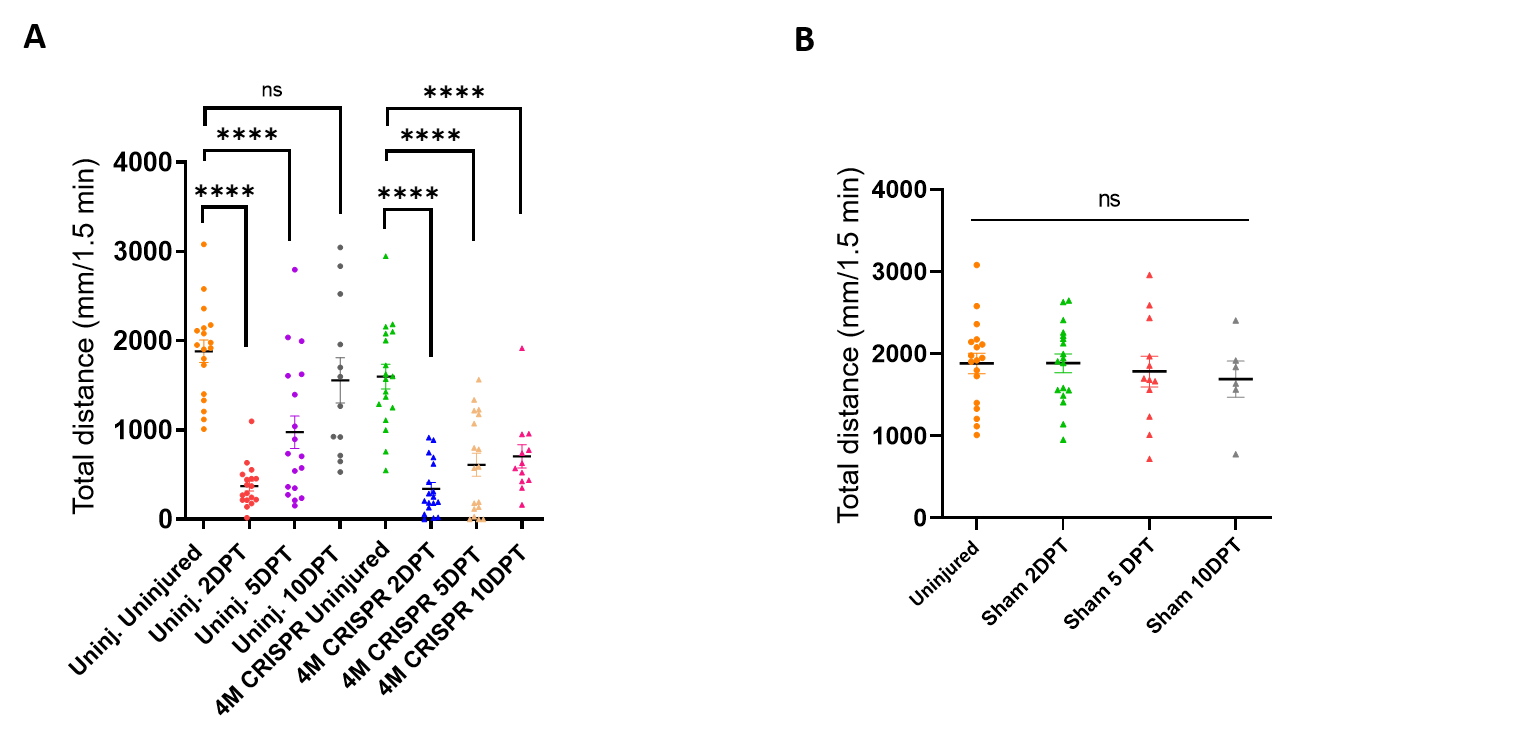

**Fig. S16: Functional recovery after spinal cord transection is compromised after knockout of Marcks and Marcksl1.**

A: Swimming recovery test in stage 50 wild-type and CRISPR tadpoles after spinal cord injury. The graph shows the total swimming distance in uninjected and 4M CRISPR-injected animals before the injury and at 2-, 5-, and 10- DPT. The y-axis indicates the distance traveled by the tadpoles in millimeters for 90 seconds (same data as in Fig. 6A but different statistical comparison). Significance was determined using an ordinary one-way ANOVA with uncorrected Fisher’s LSD for multiple comparisons. NS, not significant; ***P<0.001. All data points are shown with the line at mean. Error bars represent SEM. B: Swimming test in stage 50 sham-operated tadpoles reveals no effect on swimming performance. The graph shows the total swimming distance before the injury and 2-, 5-, and 10-days post-sham-injury (DPI). Significance was determined using an ordinary one-way ANOVA with uncorrected Fisher’s LSD for multiple comparisons. NS is not significant. All data points are shown with the line at mean. Error bars represent SEM.

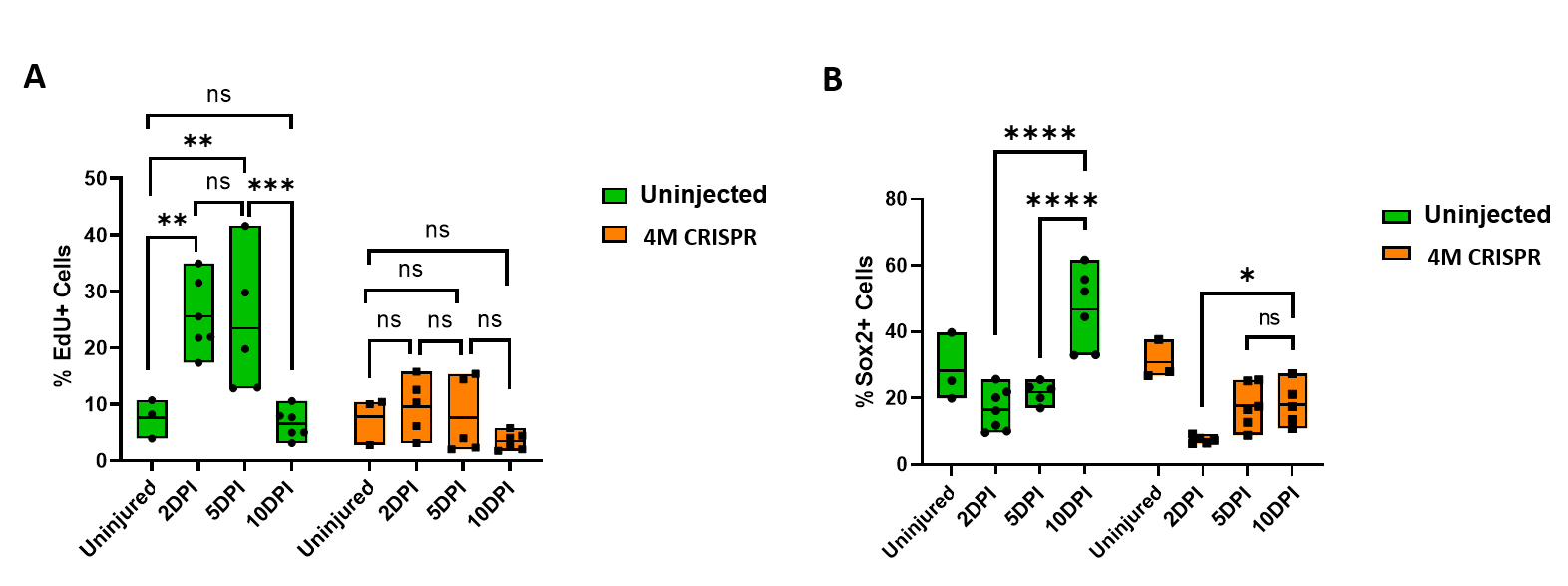

**Fig. S17: Upregulation of cell proliferation and Sox2+ neuro-glial progenitors after spinal cord transection is compromised after the knockout of Marcks and Marcksl1.**

A: Quantification of EdU+ cells at 2-, 5-, and 10-DPI in sham-operated tadpoles at stage 50 (same data as in Fig. 7 I but different statistical comparison). B: Quantification of Sox2+ cells at 2-, 5-, and 10-days post transection (DPI) in sham-operated tadpoles at stage 50 (same data as in Fig. 8 I but different statistical comparison). Significance was determined using a 2-way ANOVA with Sidak’s multiple comparisons test. NS, not significant; *P<0.05; **P≤0.01; ***P<0.001; ****P<0.0001. All data points are shown with the line at mean.

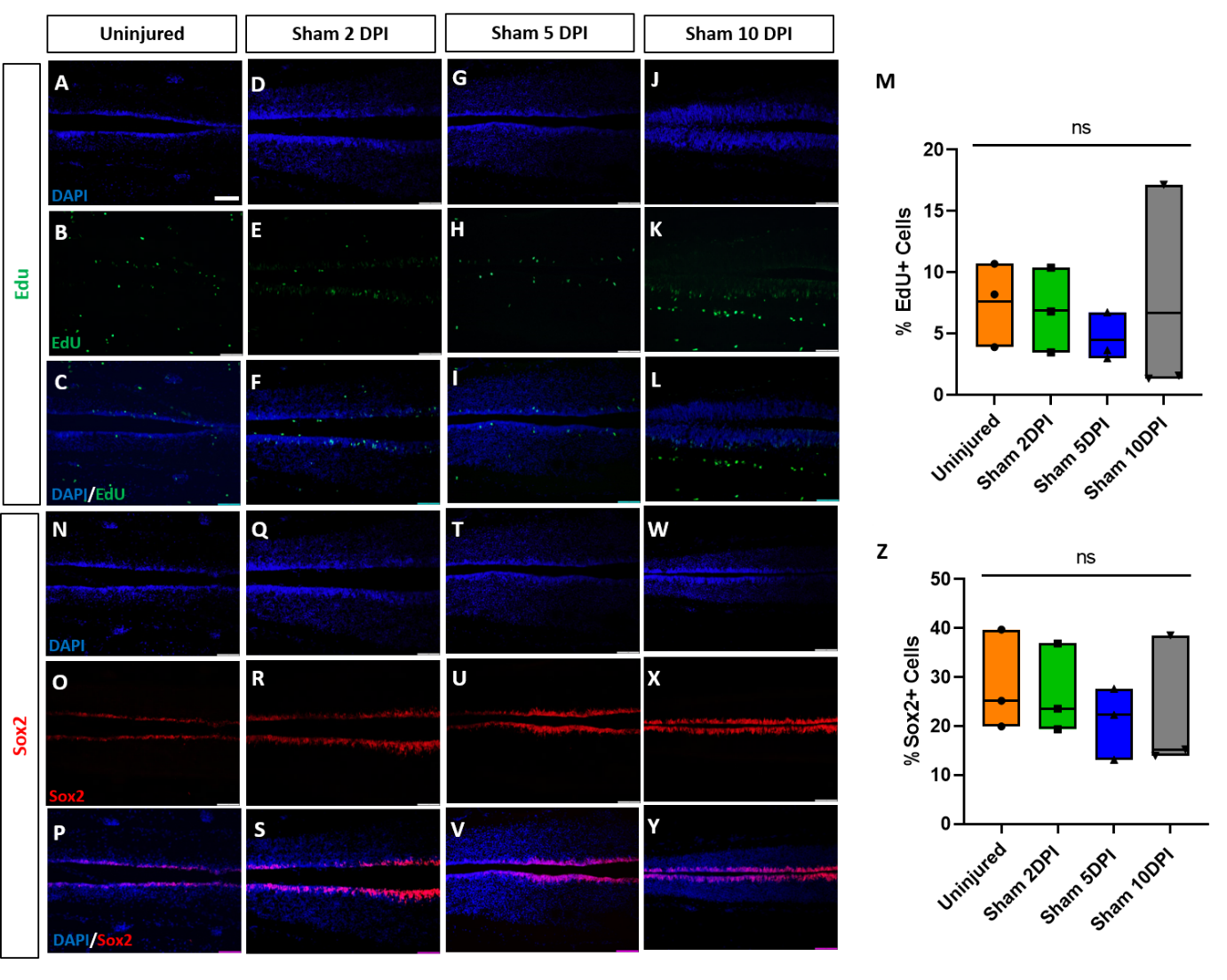

**Fig. S18:** **Sham-operated animals show no alterations of proliferation or numbers of neuro-glial progenitors.**

Evaluation and quantification of EdU+ proliferative cells (A-M) and Sox2+ neuro-glial progenitors (N-Z) at 2-, 5-, and 10-DPI in horizontal sections of sham-operated tadpoles at stage 50. Significance was determined using a 2-way ANOVA with Sidak’s multiple comparisons test. NS, not significant; *P<0.05; **P≤0.01; ***P<0.001; ****P<0.0001. All data points are shown with the line at mean. Scale bar in A= 50 µm (for all panels).
